# Integration of *In Vitro* and *In Silico* Approaches Enables Prediction of Drug-Induced Liver Injury

**DOI:** 10.1101/2025.10.24.684359

**Authors:** René Geci, Ahenk Zeynep Sayin, Stephan Schaller, Lars Kuepfer

## Abstract

Drug-induced liver injury (DILI) is a major cause of drug attrition and poses a significant threat to patient safety. However, current preclinical prediction methods, including heuristic screening rules, *in vitro* assays, machine learning models and animal testing, have serious limitations. Here, we demonstrate that combining *in vitro* toxicity data (cytotoxicity, mitochondrial toxicity, bile salt export pump (BSEP) inhibition) with pharmacokinetic information enables high DILI predictivity. In a retrospective analysis of 241 drugs, we show that the ratio of their *in vivo* maximum plasma concentration (Cmax) to their lowest *in vitro* toxicity strongly correlates with clinical DILI risks, with ROC AUC up to 96%. Then, we show that comparable predictivity (ROC AUC up to 91%) is achievable prospectively when Cmax values are predicted *in silico* by high-throughput physiologically based kinetic modelling. Dynamic simulations of bile acid perturbations further identify drugs potentially causing DILI specifically through BSEP inhibition, providing additional mechanistic insights. This integrative, mechanistic approach shows enhanced DILI predictivity and interpretability, offering an animal-free alternative for early drug development.

## Introduction

Drug-induced liver injury (DILI) remains a leading cause of drug development failures, resulting in estimated losses of billions of dollars to the pharmaceutical industry and contributing to many patient deaths each year (Watkins 2011; Hosack et al. 2023). Despite advances in toxicological methods, the ability to accurately predict DILI risks of drugs at the preclinical stage remains limited. In clinical practice, DILI can present in different patterns, as hepatocellular, cholestatic, or mixed injury, and often manifests idiosyncratically without clear dose dependence (Hoofnagle and Björnsson 2019). This makes it difficult to detect in preclinical toxicity screenings (Kullak-Ublick et al. 2017). Given its substantial impact on both drug attrition and patient safety, improving early-stage DILI prediction remains a critical unmet need.

Animal testing has been the historical gold standard for predicting drug toxicity in humans. However, reported concordance rates between animal and human liver toxicity outcomes are low, ranging from 40% to 70% (Olson et al. 2000; Fourches et al. 2010; Leenaars et al. 2019), depending on the dataset and endpoint. Additionally, there are growing ethical concerns and societal pressure on policymakers and regulators to phase out animal testing. This further underscores the need for new human-relevant toxicity assessment methods.

Early preclinical efforts to assess DILI risks of drug candidates often rely on simple heuristic screening rules using readily available drug properties, such as lipophilicity or dose (Chen et al. 2013). While easy to apply, those rules lack a mechanistic basis and frequently misclassify compounds. To address these limitations, various *in vitro* test assays have been developed which measure drug interactions with relevant toxicity endpoints, such as cytotoxicity, mitochondrial dysfunction, or bile salt export pump (BSEP) inhibition (Weaver et al. 2020; Yucha et al. 2017). Such assays are nowadays applied in standard preclinical safety workflows (Walker et al. 2020; Norman 2020). However, their toxicity readouts are typically analysed without consideration of pharmacokinetic exposure information, and their predictivity often remains limited (Dirven et al. 2021; Taylor et al. 2025).

A succinct example of the challenges for using *in vitro* toxicity methods to predict DILI is the measurement of BSEP inhibition. BSEP is a well-characterised hepatic transporter involved in bile acid metabolism and has been mechanistically linked to cholestatic forms of DILI (Dawson et al. 2012; Stieger 2010; Vinken et al. 2013). Nonetheless, BSEP inhibition data alone have shown inconsistent performance for predicting clinical DILI outcomes and, in some cases, perform no better than heuristic screening rules (Chan and Benet 2018a). This poor predictivity has led to the questioning of the utility of *in vitro* assays, such as BSEP inhibition testing, with some authors dismissing their predictive utility entirely (Chan and Benet 2018b). For mechanisms less well-characterised than BSEP inhibition, *in vitro* to *in vivo* translatability is even more challenging.

Another now frequently explored strategy for DILI prediction involves the use of machine learning (ML) approaches trained on chemical descriptors, *in vitro* assay and/or *in vivo* toxicity data (Seal et al. 2024b; Garcia de Lomana et al. 2025; Minerali et al. 2020; Williams et al. 2020). But while some ML models show promising performances, most still operate as black boxes without mechanistic transparency (Jia et al. 2023; Hemmerich and Ecker 2020). This limits their interpretability, prospective use and further poses challenges for regulatory acceptance. Additionally, the ability of such models to extrapolate predictions to structurally novel compound classes often remains uncertain.

In summary, none of the current toxicity assessment approaches have so far been able to provide the predictivity of DILI risks needed for efficient drug discovery, nor do they offer the transparency and mechanistic interpretability required by regulators seeking alternatives to animal testing. In this study, we demonstrate that accurate DILI risk predictions are achievable by integrating already available *in vitro* toxicity assays with *in vivo* pharmacokinetic (PK) information. Furthermore, even fully prospective DILI risk assessments of novel drug candidates become possible by using a recently developed high-throughput physiologically based kinetic (HT-PBK) modelling approach to predict PK parameters *in silico*.

## Methods

### Data collection

We searched the literature for studies that had published *in vitro* hepatotoxicity measurements to predict the DILI risks of drugs, and which had determined specific toxicity potency values, e.g., IC50 concentrations, instead of binary active/non-active hit readouts. We then integrated all these measurements into a single dataset by matching the various compounds against each other using their canonical SMILES identifiers retrieved from PubChem (Kim et al. 2025). Salt ions were removed and matching was performed based on the presumed active pharmaceutical ingredient. Eventually we collected *in vitro* hepatotoxicity data from seventeen *in vitro* hepatotoxicity datasets (Aleo et al. 2020; Shah et al. 2015; Morgan et al. 2013; Dawson et al. 2012; Warner et al. 2012; Fäs et al. 2024; Ewald et al. 2025; Albrecht et al. 2025; Proctor et al. 2017; Braak et al. 2024; O’Brien et al. 2006; Gustafsson et al. 2014; Persson et al. 2013; Schadt et al. 2015; Williams et al. 2020; Porceddu et al. 2012; U.S. Environmental Protection Agency 2025). Additionally, we matched compounds against the FDA-approved DILIrank (Chen et al. 2016) database to retrieve classifications of their clinical DILI risk. Some of the retrieved studies already included data on the *in vivo* Cmax values of drugs, however, those did not always match each other. Therefore, we manually retrieved information about the highest recommended single oral dose and *in vivo* Cmax values of corresponding clinical studies for every drug. Drugs for which no clinical DILI risk or no single oral dose Cmax could be identified were removed from the dataset. When multiple *in vitro* potency values had been measured for the same functional toxicity mechanism (cytotoxicity, mitochondrial toxicity), then those values were integrated by using the lowest (strongest potency) value, assuming that the assays and cell systems used may have had varying abilities to detect perturbations of the underlying endpoint. When multiple BSEP IC50 values were available, those were integrated by using the median value, assuming that BSEP inhibition assays covered the same endpoint with the same sensitivity and that differences between those measurements were technical variability.

To analyse in detail which drugs showed a specifically cholestatic pattern of DILI, we had to generate our own database, as we were not aware of any standardised databases comparable to DILIrank for drug-induced cholestasis. For this, we integrated cholestasis classifications from the adverse drug reactions’ database SIDER (Kuhn et al. 2016) and publications which stated drugs’ DILI patterns (Kotsampasakou and Ecker 2017; Lima Toccafondo Vieira and Tagliati 2014; Riede et al. 2017; van Brantegem et al. 2020; Bruijn and Rietjens 2024; Drees et al. 2025). If drugs were referenced in one of those sources to have cholestatic or hepatocellular DILI patterns, or to be safe, or if multiple sources were all in agreement with each other regarding the same compound, then those classifications were adopted. However, if different sources classified the same compound differently, we labelled those drugs as “contradiction”. Compounds that failed in clinical development were labelled as such, since they could not be expected to be represented in any of the used resources. All other compounds, for which no DILI pattern information could be matched were labelled “unknown”.

### High-throughput PBK simulations

High-throughput physiologically based kinetic (HT-PBK) simulations of drugs were based on previously performed model validations (Geci et al. 2024; Gadaleta et al. 2024). Briefly, all drug properties required for PBK modelling (Kuepfer et al. 2016) were predicted using previously evaluated *in silico* tools: ADMETLab LogD, LogS, fraction unbound, CACO2 permeability (Fu et al. 2024), ADMET Predictor v12 Fasted State Simulated Intestinal Fluid solubility (https://www.simulations-plus.com/software/admetpredictor), and PKsmart clearance (Seal et al. 2024a). For each compound, the default whole-body PBK model implemented in PK-Sim (Willmann et al. 2003) version 11.1.137 was parameterised from R version 4.2.2 (R Core Team 2022), and oral administration of the extracted highest oral dose was simulated. For partitioning, the method PK-Sim Standard (Willmann et al. 2005) was used and as readouts simulated Cmax values were extracted. To account for the fact that dissolution times of the different drug formulations were unknown in the retrospective analysis, while they would be known for prospective evaluations of a specific drug formulation, we performed simulations with different Lint80 dissolution times (1-180 min) and eventually used simulated Cmax values of drugs closest to their observed values.

### Bile acid model simulations

To simulate the dynamic effects of BSEP inhibition by drugs, and subsequent intrahepatic bile acid changes, we used a previously reported Physiologically based bile acid (PBBA) model (Kister et al. 2025). We implemented drug interactions with BSEP as competitive inhibition in the BSEP-mediated transport rate as

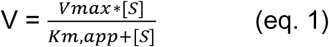

with

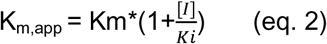

Where V is the velocity of the BSEP-mediated transport, [S] is the bile acid substrate concentration, V_max_ is the maximal BSEP transport velocity, [I] is the inhibitor concentration and K_m_ the Michaelis-Menten constant of the transported bile acid substrates.

To parameterise eq. 2, we converted measured IC50 values of drugs to Ki values using the equation (Cheng and Prusoff 1973)

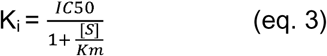

with K_i_ being the inhibition constant of drugs, IC50 the measured half-maximal inhibition concentration and K_m_ the Michaelis-Menten constant of the substrates used as *in vitro* probes. To dynamically represent drug concentrations, we used the plasma concentration-times profiles of drugs simulated using the HT-PBK method. Since bile acid model simulations were used for retrospective analysis, *in vivo* observed (Lombardo et al. 2024) plasma clearances instead of predicted values were used, if available. As readouts of the bile acid model, we extracted the concentration of intrahepatic total conjugated bile acid concentration and determined the duration of elevation of this concentration above two-fold, relative to the unperturbed state, and the duration of elevation to more than 10% above the unperturbed peak concentration.

## Results

### Integrated hepatotoxicity dataset overview

After integrating seventeen *in vitro* hepatotoxicity datasets, we had obtained *in vitro* potency measurement results of 487 unique compounds. However, only for 355 of them a specific toxicity potency value had been reported, while for the others, even at the highest test concentration, no measurable effect was observed. We searched for information on the highest recommended oral dose and corresponding maximum plasma concentration (Cmax) values of these 355 active compounds, as well as on their clinical DILI outcomes using DILIrank (Chen et al. 2016). Eventually, we ended up with a complete, integrated dataset of 241 unique drugs of which the oral dose, corresponding Cmax, clinical DILI outcome and at least one *in vitro* toxicity value were available (Fig. 1).

**Fig. 1:**
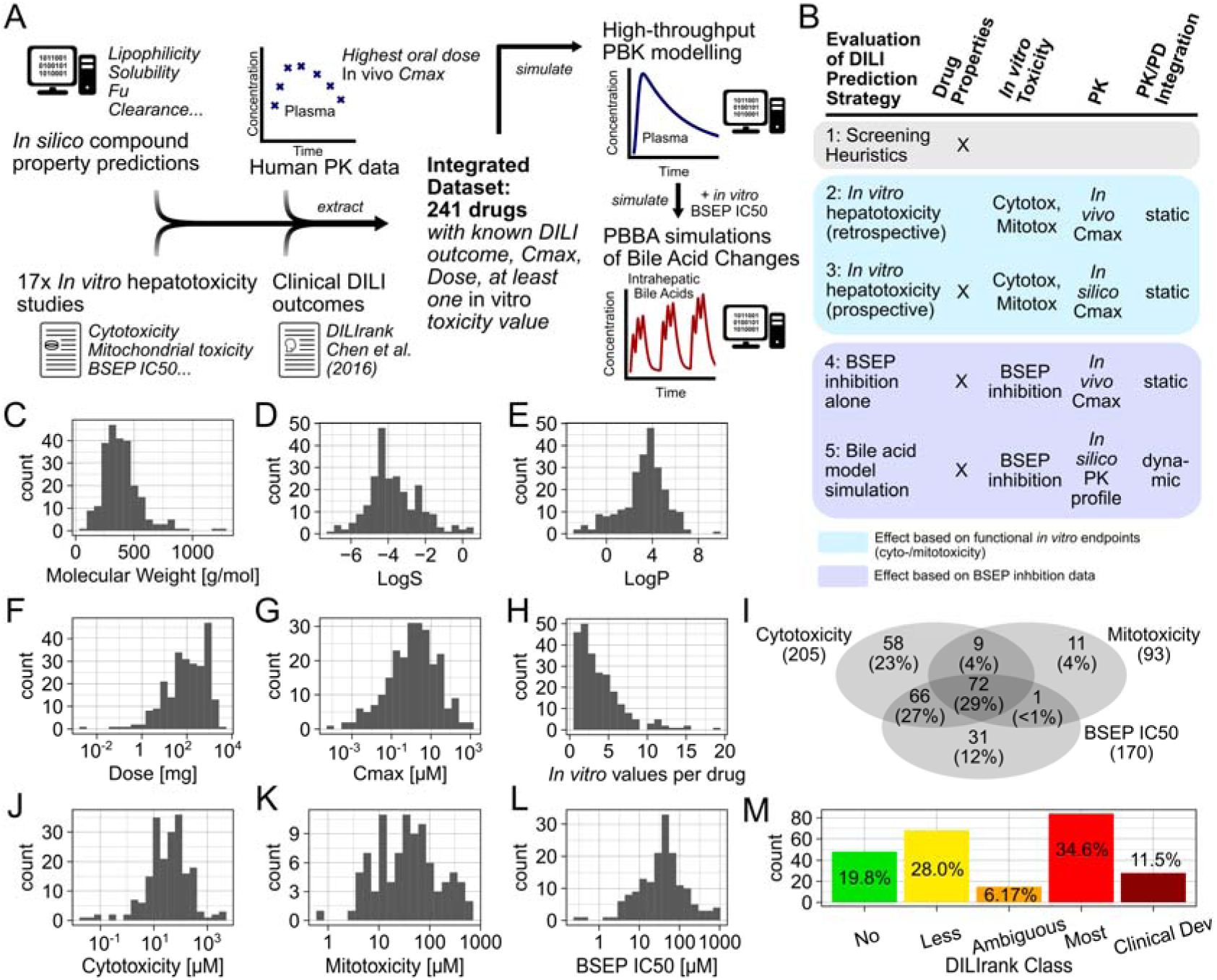
Data collection, analysis workflow and drug properties. Various literature datasets were integrated to obtain a dataset of 241 drugs with complete information on *in vitro* hepatotoxicity, clinical DILI outcome and pharmacokinetics (A). DILI prediction strategies of varying complexity were evaluated using the integrated dataset (B). Statistical plots of various drug properties and parameters of the collected dataset (C-M).

The most frequent mechanism of *in vitro* toxicity available in our dataset was cytotoxicity (205/241), which had been measured in a variety of cell systems across different studies, followed by BSEP inhibition (170/241) and mitochondrial toxicity (93/241). Some compounds had been tested in multiple *in vitro* studies and had up to 19 unique potency values available, whereas others had only been tested in one assay. The drugs of the integrated dataset represented all DILIrank concern categories, with the majority being Most-(34.6%), Less-DILI concern (28.0%), and Clinical Development Failures (11.5%, as assigned previously (Aleo et al. 2020)). This high representation of DILI-positive compounds was due to fewer DILI-negative compounds having been tested in the *in vitro* studies, and because many of those negative control compounds did not yield an *in vitro* measurable toxicity activity response. Nonetheless, our final dataset included 19.8% No-DILI concern compounds with at least one *in vitro* potency value determined. The 241 compounds of the final dataset represented a diverse set of small molecules across different molecular properties (Fig. 1C-M). As the hepatotoxicity potential of Ambiguous-DILI concern drugs remains clinically inconclusive, we restricted our analysis to drugs of the remaining conclusive DILIrank categories.

### Evaluation 1: Simple heuristic screening rules poorly predict DILI

Many heuristic screening rules have been proposed for the pragmatic prediction of DILI at the preclinical stage, for example based on the lipophilicity or the dose of a drug. Since such heuristics are used in the pharmaceutical industry, and as a benchmark for later evaluations, we first analysed how well those pragmatic rules would predict DILI-positive compounds of our integrated dataset. The first often used heuristic states that compounds with higher LogP are more likely to be DILI-positive (Chen et al. 2013). However, when comparing the LogP values of compounds, it appeared that LogP alone did not allow for a meaningful distinction of the different DILIrank classes (Fig. 2A). The second frequently used heuristic for DILI prediction states that the dose of a drug is a DILI risk factor (Lammert et al. 2008), assuming that higher doses are related to higher DILI risks. For drug doses, we did find larger differences between DILI rank classes, with No-DILI concern compounds having the lowest dose values, and Most-DILI compounds having the highest doses (Fig. 2B). LogP and dose have also been combined in the “Rule of Two” (Chen et al. 2013), which resulted in a higher specificity of DILI predictions than when using each property on its own, although at the cost of lower sensitivity (Fig. 2E; SI-Fig. 1).

**Fig. 2:**
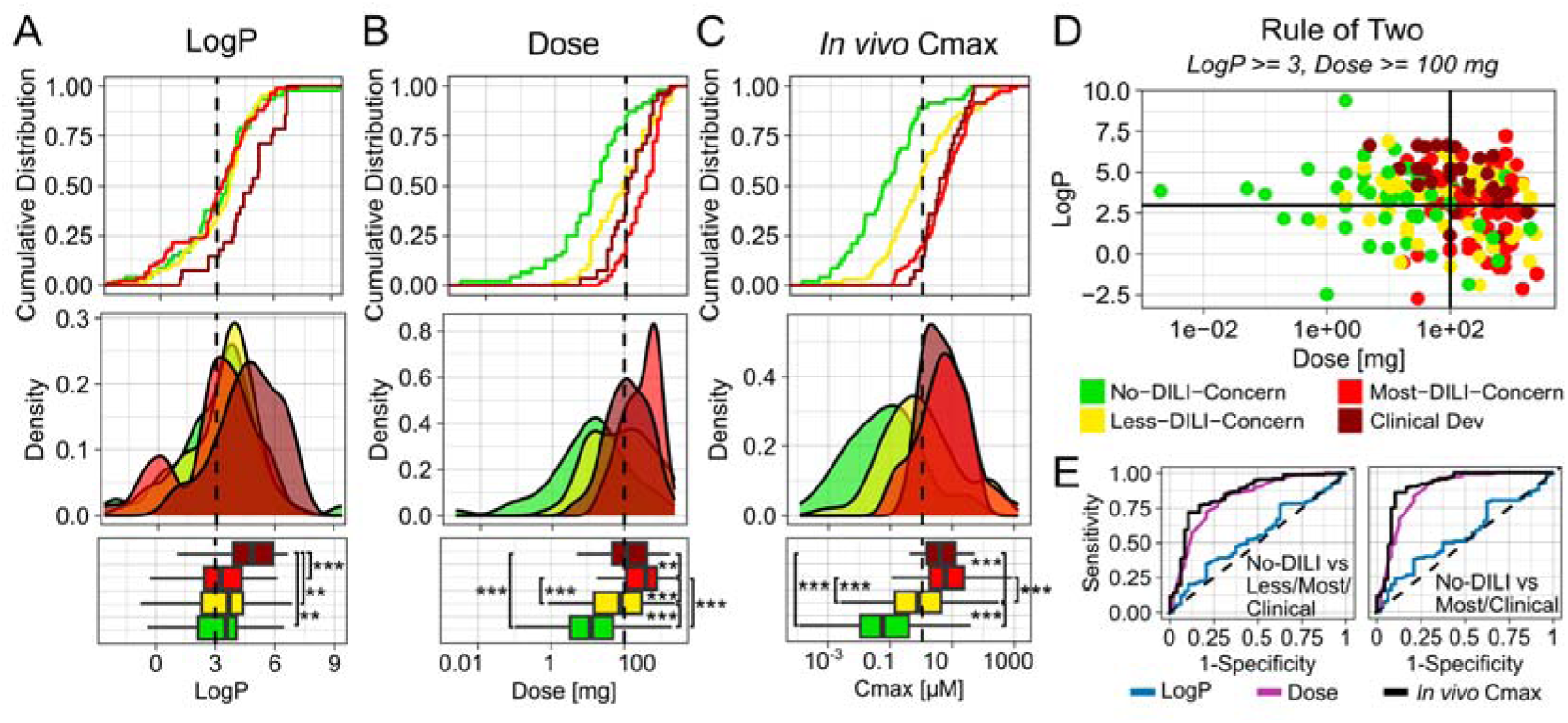
Predictivity of simple DILI screening heuristics (evaluation 1). Cumulative distribution, probability density and boxplots of properties used for heuristic drug candidate screening, like LogP (A), Dose (B) and *in vivo* Cmax (C) for drugs of the different DILIrank classes. LogP values were predicted using ADMETLab. Distribution of drugs across the Rule of Two quadrants (D). ROC curves of the different DILI screening heuristics for distinguishing No-DILI compounds from Less-/Most-DILI/Clinical Development Failures (Aleo et al. 2020), or only from Most-DILI/Clinical Development Failures (E). Boxplots show the median and interquartile range; whiskers extend to 1.5×IQR. Outliers were excluded from the plot for clarity but were included in statistical analyses. Statistical comparisons on log-transformed values were performed using pairwise Wilcoxon rank-sum tests with Holm correction: ** p < 0.01, *** p < 0.001.

Finally, the Cmax values of drugs have been proposed as a predictor of DILI risk, and in line with previous reports (Shah et al. 2015), we found that Cmax performs better than any other heuristic. *In vivo* Cmax on its own enabled the discrimination between No-from Most-DILI and Clinical Development Failure compounds with a specificity of 89.6% and a sensitivity of 82.1%. The observation that Cmax performed better than dose was in line with our expectations, since dose presumably only serves as a surrogate for the *in vivo* concentrations of drugs, and Cmax is a more direct reflection of that. But such high predictivity was still unexpected, as it suggested that the toxic potency of drugs may be barely relevant for clinical DILI outcomes. From a mechanistic perspective, one would assume that both *in vivo* concentration and toxic potency contribute equally to *in vivo* toxicity outcomes, and extensive efforts have focused on developing *in vitro* systems to test for this property. But all collected *in vitro* toxicity values of drugs showed little ability for distinguishing DILI concern classes on their own (SI-Fig. 1). Even when *in vitro* toxicity values were integrated into a single lowest potency value per compound, those lowest potency values performed better than each of the *in vitro* values individually, but still not better than simple heuristic rules (SI-Table 1).

### Evaluation 2: Cmax to in vitro toxicity ratios strongly predict DILI

Despite the observation that *in vitro* toxicity data alone did not perform better than simple heuristics for distinguishing drugs of different DILIrank classes, we next tested whether they would contribute additional signal for DILI predictions when being combined with pharmacokinetic information (*in vivo* Cmax values). Such exposure-activity integrations are increasingly employed in toxicology (Wetmore et al. 2015; Middleton et al. 2022; Cable et al. 2024), but for DILI predictions, they so far only showed mixed success (Morgan et al. 2013; Shah et al. 2015). In several studies, their predictive accuracy did not clearly surpass that of simple property-based heuristics, prompting debate over the practical utility of this mechanistic strategy. For the first evaluation, we exclusively relied on data from functional *in vitro* toxicity endpoints (cytotoxicity and mitochondrial toxicity), initially leaving out BSEP inhibition data.

When calculating the ratio of *in vivo* Cmax to lowest functional *in vitro* toxicity of each compound, we found that those ratios correlated well with clinical DILI outcomes (Fig. 3A). The ratios of DILI concern classes were significantly different from each other (Fig. 3C) and values aligned with the severity of DILIrank classifications. No-DILI compounds possessed the lowest ratio values, Less-DILI showed intermediate, and Most-DILI compounds showed the highest ratios. The AUC of the ROC curve for this ratio was 96% for distinguishing No-from Most-DILI and Clinical Development Failure drugs, and 90% for separating No-from Less-, Most-DILI and Clinical Development Failures (Fig. 3D). The best threshold value for classification as determined by Youden’s index (YOUDEN 1950) was 1.1%, with a specificity of 82% and sensitivity of 97% for distinguishing No-from Most-DILI and Clinical Development Failure compounds.

**Fig. 3:**
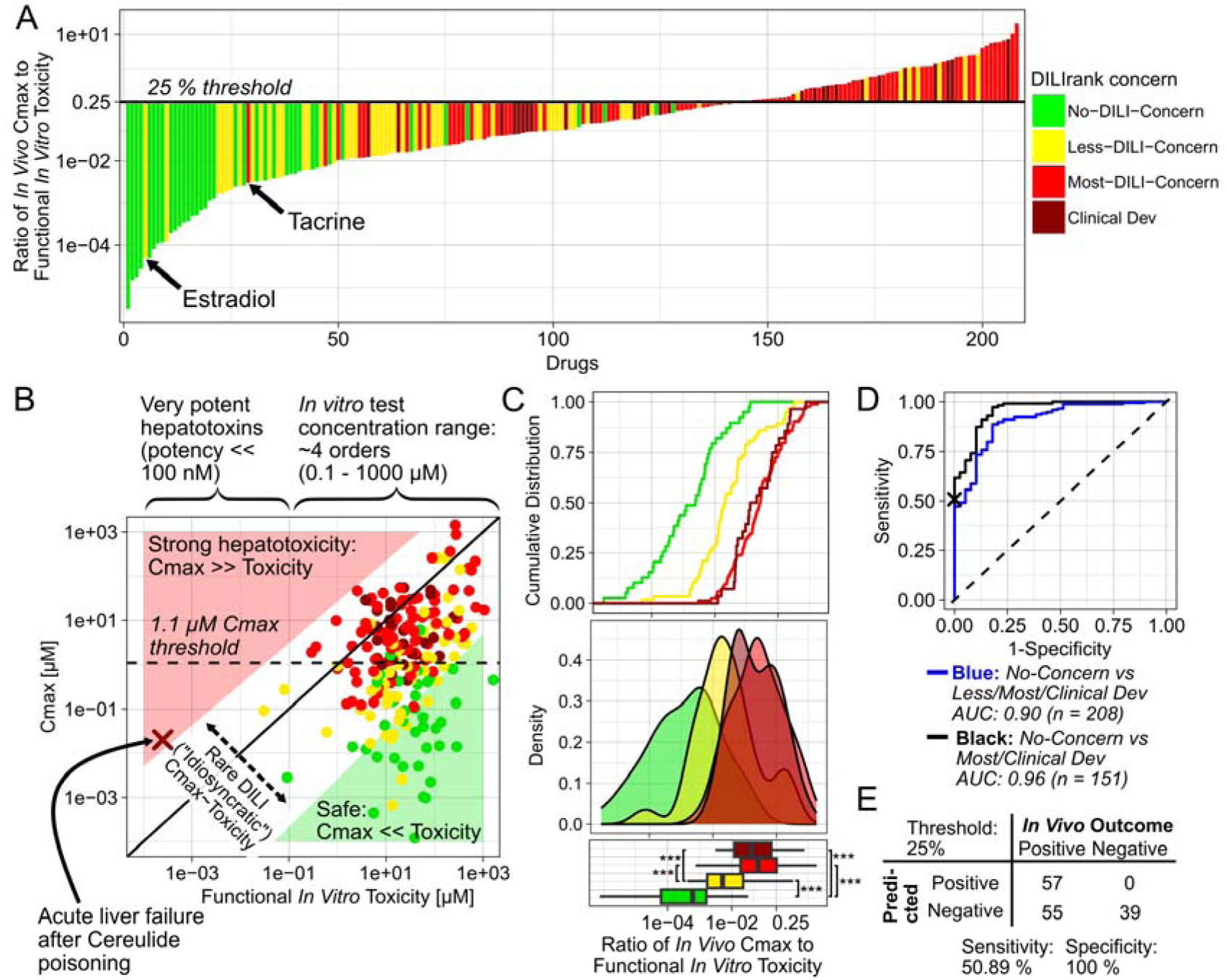
DILI predictivity of integrated functional *in vitro* toxicity and *in vivo* Cmax data (evaluation 2). Ratio of *in vivo* Cmax to lowest functional *in vitro* toxicity of each drug (A). Functional toxicity refers to all *in vitro* toxicity data except BSEP inhibition. *In vivo* Cmax values against lowest functional *in vitro* toxicity values of each drug (B). The red cross indicates a reported case of liver failure after cereulide poisoning. Green and red triangles mark areas where *in vivo* Cmax is much smaller or larger than functional *in vitro* toxicity, respectively. Cumulative distribution, probability distribution and boxplots of the ratio of *in vivo* Cmax to functional *in vitro* toxicity values of different DILIrank classes (C). Receiver operating characteristic curves of *in vivo* Cmax to toxic potency ratios for distinguishing No-DILI compounds from DILI-positive drugs (D). Boxplots show the median and interquartile range; whiskers extend to 1.5×IQR. Outliers were excluded from the plot for clarity but were included in statistical analyses. Statistical comparisons on log-transformed values were performed using pairwise Wilcoxon rank-sum tests with Holm correction: ** p < 0.01, *** p < 0.001. Predictive performance parameters of ratios using 25% as threshold for DILI positive classification (E).

However, from a mechanistic perspective, an *in vivo* concentration that is only 1.1% of the *in vitro* potency seemed too low that it could explain biologically meaningful perturbations *in vivo*. Instead, we chose 25% as a mechanistically more plausible threshold for the lowest limit of concentrations required for *in vivo* toxicity, which resulted in a specificity of 100% and sensitivity of 51%. In other words, this mechanistic threshold yielded no false-positive but some false-negative predictions, corresponding to compounds being clinically DILI-positive but with Cmax to *in vitro* potency ratios below 25%.

To understand why some compounds were falsely predicted to be DILI-negative, we investigated them in more detail. Estradiol, classified as Less-DILI concern, was the DILI-positive compound presenting the lowest Cmax to *in vitro* toxicity ratio (4.9x10^-5^) and is an endogenous sex hormone used in hormone replacement and gender-affirming therapy. After oral administration of 2 mg, its *in vivo* Cmax is in the sub-nanomolar range (0.1 -0.5 nM), whereas its eight *in vitro* toxicity values range from 13.5 to 103 µM, hence more than 1000-fold higher than its Cmax. Given estradiol’s endogenous nature, it is unsurprising that it would not cause cytotoxicity or mitochondrial toxicity *in vivo*. One potential explanation for its observed hepatotoxicity is that it causes DILI through other mechanisms than general cytotoxicity or mitochondrial toxicity. In fact, the deregulation of bile acid (BA) synthetic or metabolic enzymes, and BA transporters have been discussed as key mechanisms for its DILI potential (Zu et al. 2021). But such gene expression perturbations were not captured by the *in vitro* systems included here. In animal models, its toxicity has further been linked to the generation of conjugated metabolites like estradiol-17β-glucuronide (Meyers et al. 1980), which may not be adequately generated *in vitro* either.

The Most-DILI concern drug with the lowest Cmax to *in vitro* toxicity ratio was tacrine (3x10^-3^). LiverTox (2020) states that its precise mechanism of liver injury is unknown, but that it is extensively metabolised and that the production of a toxic intermediate likely plays a central role in its toxicity, with intestinal bacteria possibly being an important factor (Yip et al. 2018). Again, these mechanisms fall outside the scope of the *in vitro* hepatotoxicity assays included here. Based on the manual review of false negatives, we concluded that, despite its lower sensitivity, the mechanistic classification threshold value of 25% seemed more appropriate for further evaluations than the statistically more optimal lower threshold of 1.1%.

When directly comparing *in vivo* Cmax values of drugs to their *in vitro* toxicity (Fig. 3B), we further observed that the ranges of those values differed substantially. While Cmax values were distributed over 7 orders of magnitude (10^-4^ to 10^3^ µM), the *in vitro* toxicity values only varied across 4 orders (10^-1^ to 10^3^ µM). The latter can partially be explained by the fact that this reflects the concentration ranges used for *in vitro* testing in the studies of which we retrieved data. However, this revealed a bias in our integrated dataset, and similarly in previously performed analyses using those data, which explains why dose or Cmax on their own may often have been observed to be such powerful predictors of DILI. If all analysed compounds have relatively similar toxic potency, then the main factor driving differences in their clinical toxicity outcomes could only be their *in vivo* concentration. This apparent predictive power of dose or Cmax alone could only have been challenged by including compounds showing clinical toxicity at low concentrations, which would have required testing highly potent hepatotoxins, or compounds that, due to very low toxic potency, do not cause clinical toxicity even at high concentrations, like endogenous metabolites or natural compounds. But both groups of substances were not included in the *in vitro* studies. Even if they had been, their extremely high or low potency would not have been determinable in the fixed *in vitro* test concentration ranges. To illustrate that compounds with low dose or Cmax values are not generally safe and can still cause even severe hepatotoxicity, we added a case report of liver failure after poisoning with cereulide to Fig. 3B (Schreiber et al. 2021; Decleer et al. 2018).

Furthermore, our analysis offered an explanation why DILI may typically be observed only in an “idiosyncratic” manner, without obvious dose dependence. None of the drugs in our analysis possessed *in vivo* Cmax values substantially exceeding their corresponding *in vitro* toxicity, not even most compounds classified as Most-DILI concern. This is plausible, since otherwise these drugs would presumably show very clear *in vivo* hepatotoxicity, and then they would never have been approved by the FDA in the first place. But if the Cmax concentrations of even those DILI-positive drugs, at their highest recommended dose, are still typically below their toxic potency, then this may explain why these drugs are still safe in most patients, and why clinically DILI toxicity is only observed “idiosyncratically” in the most susceptible subgroups of patients, like those with genetic predispositions, comorbidities, comedications etc. In pharmacology, it is usually attempted to achieve *in vivo* concentrations multiplelZlfold above the *in vitro* potency of a drug to ensure strong predictable *in vivo* effects (Maurer et al. 2020). However, we found that almost none of the DILI-positive drugs fulfilled this condition.

### Evaluation 3: In silico predicted Cmax enables prospective DILI evaluation

Retrospective analysis combining functional *in vitro* toxicity data with *in vivo* observed Cmax values revealed that *in vitro* assays provide useful information for the prediction of DILI and that combining both data types allowed good predictions of clinical DILI risks. However, it is only possible to use *in vivo* observed Cmax values for retrospective analyses, once human clinical studies have been performed already. A key limiting factor for prospective DILI predictions of new drug candidates has so far been the unavailability of Cmax values before any clinical studies have been performed. Recently, we developed a high-throughput method that integrates machine learning models with mechanistic PBK modelling to predict human pharmacokinetics, and thereby enables the prospective, *in silico*-based prediction of drug Cmax values.

We applied this method to our integrated dataset and compared *in silico*-predicted Cmax values of drugs against *in vivo* observed values (Fig 4C). As reported previously, we observed that most (90%) drugs’ Cmax values were predicted within the 10-fold range. Then, we used those predicted values, instead of *in vivo* observed ones, for calculating the Cmax to *in vitro* toxicity ratios of drugs (Fig. 4A). Unlike when using *in vivo* Cmax, the use of *in silico-*predicted Cmax values did result in some false-positive DILI risk predictions. The two No-DILI compounds phenazopyridine and etidronate were predicted to have higher Cmax values than they exhibited *in vivo,* and subsequently also to possess higher Cmax to *in vitro* toxicity ratios. But, overall, predicted Cmax to *in vitro* toxicity ratios still correlated well with DILIrank labels. ROC AUC values were only slightly lower than when using *in vivo* Cmax values (86% instead of 90%, or 91% instead of 96%; Fig 4D). Notably, *in silico*-predicted Cmax values alone also performed better for predicting DILI risks than *in vitro* toxicity values and screening heuristics, except for the observed *in vivo* Cmax (SI-Fig. 1).

**Fig. 4:**
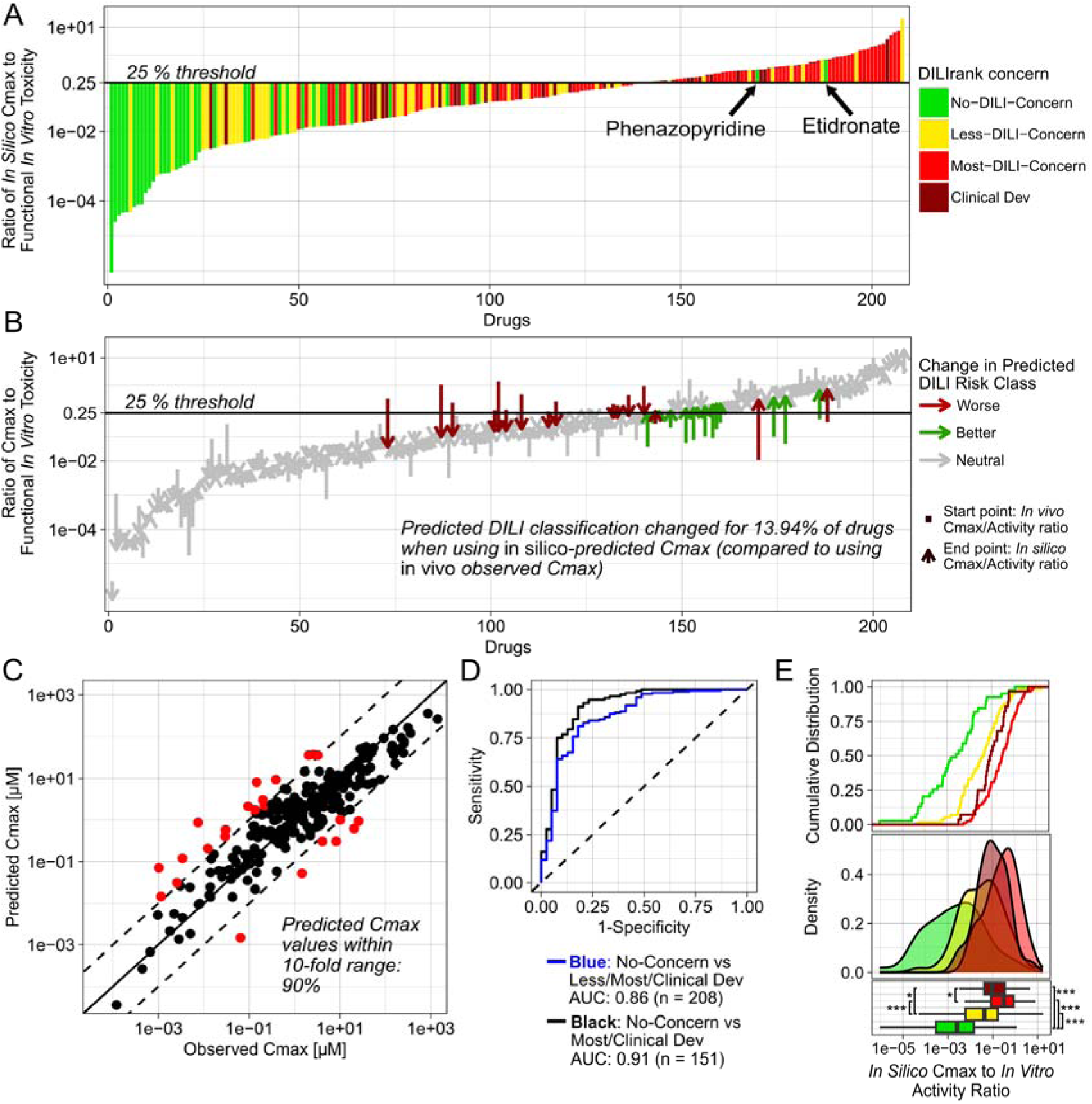
Prospective DILI predictivity of functional *in vitro* toxicity and *in silico* Cmax predictions (evaluation 3). Ratio of *in silico-*predicted Cmax to lowest functional *in vitro* toxicity values of each drug (A). Functional toxicity refers to all *in vitro* toxicity values except BSEP inhibition. Change in DILI predictions based on use of *in silico-predicted* against *in vivo* Cmax values (B). Green indicates changes to a better DILI prediction using *in silico* Cmax values compared to *in vivo* Cmax, red indicates changes to worse DILI class prediction and grey indicates no change between the two Cmax sources. Start points of arrows indicate the ratio based on *in vivo* Cmax, and end points the ratio based on *in silico-*predicted Cmax values. *In silico-*predicted values against *in vivo* observed Cmax values of drugs (C). Red indicates Cmax values outside the 10-fold range. Receiver operating characteristic curves of ratios for distinguishing No-DILI compounds from DILI-positive drugs (D). Cumulative distribution, probability distribution and boxplots of the ratio of *in vivo* Cmax to functional *in vitro* toxicity values of different DILIrank classes (E). Boxplots show the median and interquartile range; whiskers extend to 1.5×IQR. Statistical comparisons on log-transformed values were performed using pairwise Wilcoxon rank-sum tests with Holm correction: * p < 0.05, ** p < 0.01, *** p < 0.001.

As reported previously (Geci et al. 2024), we observed a tendency for *in silico*-predicted Cmax values to be overestimated, with 17 drugs being overpredicted more than ten-fold, and only 7 being underpredicted to this extent. And because DILI-positive compounds were also over-represented in our dataset, the overall predictive performance when using *in silico*-predicted Cmax values may appear inflated relative to a balanced dataset. To more truthfully assess the additional error introduced by *in silico* Cmax prediction, we quantified how often DILI classifications changed when using predicted values compared to classifications based on *in vivo* Cmax values. We found that this was the case for 29 out of 208 drugs (13.94%; Fig. 4B). While some of these changes coincidentally improved individual DILI predictions, we consider all changes as part of the additional technical error introduced by *in silico* Cmax prediction, as *in silico*-predicted values cannot be assumed to be more accurate than *in vivo* observed values.

### Evaluation 4: Inclusion of BSEP inhibition improves DILI predictivity

After evaluating the DILI predictivity of functional *in vitro* toxicity assays (cytotoxicity and mitochondrial toxicity), we next analysed the value of additionally including *in vitro* BSEP inhibition measurements. BSEP inhibition has historically been considered an important mechanism of cholestatic DILI, which is why it is a standard screening assay performed in the pharmaceutical industry. However, the relevance of *in vitro* BSEP inhibition data for the prediction of clinical DILI outcomes has been debated, so that some have claimed that “there is no support for *in vitro* BSEP inhibition being DILI predictive” (Chan and Benet 2018b).

As with functional *in vitro* toxicity data, we calculated the ratios of *in vivo* Cmax values of drugs to their BSEP IC50. Again, we found good correlation of those ratios with DILI classifications (AUC ROC up to 91%; Fig. 5). No-DILI compounds had lower ratios, and Most-DILI drugs possessed higher ratios. To ensure that the predictive power of BSEP inhibition data was not based on correlation with other functional *in vitro* activities, we investigated how the Cmax to toxicity ratios of drugs changed upon inclusion of BSEP inhibition as an additional data source (Fig. 5B). This analysis was only possible for drugs for which both BSEP IC50 and another functional *in vitro* toxicity value were available. Except for mifepristone, adding BSEP inhibition only strongly increased the ratio values of DILI-positive compounds. Notably, it appeared that the drugs most strongly affected by inclusion of BSEP inhibition were mostly Less-DILI compounds. For compounds lacking other functional *in vitro* toxicity values, either because they were not tested or because they did not show any activity, this analysis was not possible. Interestingly, all those other drugs were classified as No- or Less-DILI concern (Fig. 5C). Furthermore, when comparing the distributions of the different DILIrank classes, we observed that including BSEP inhibition appeared to particularly strongly shift the distribution of Less-DILI compounds towards larger Cmax to *in vitro* toxicity ratios (Fig. 5D-F).

**Fig. 5:**
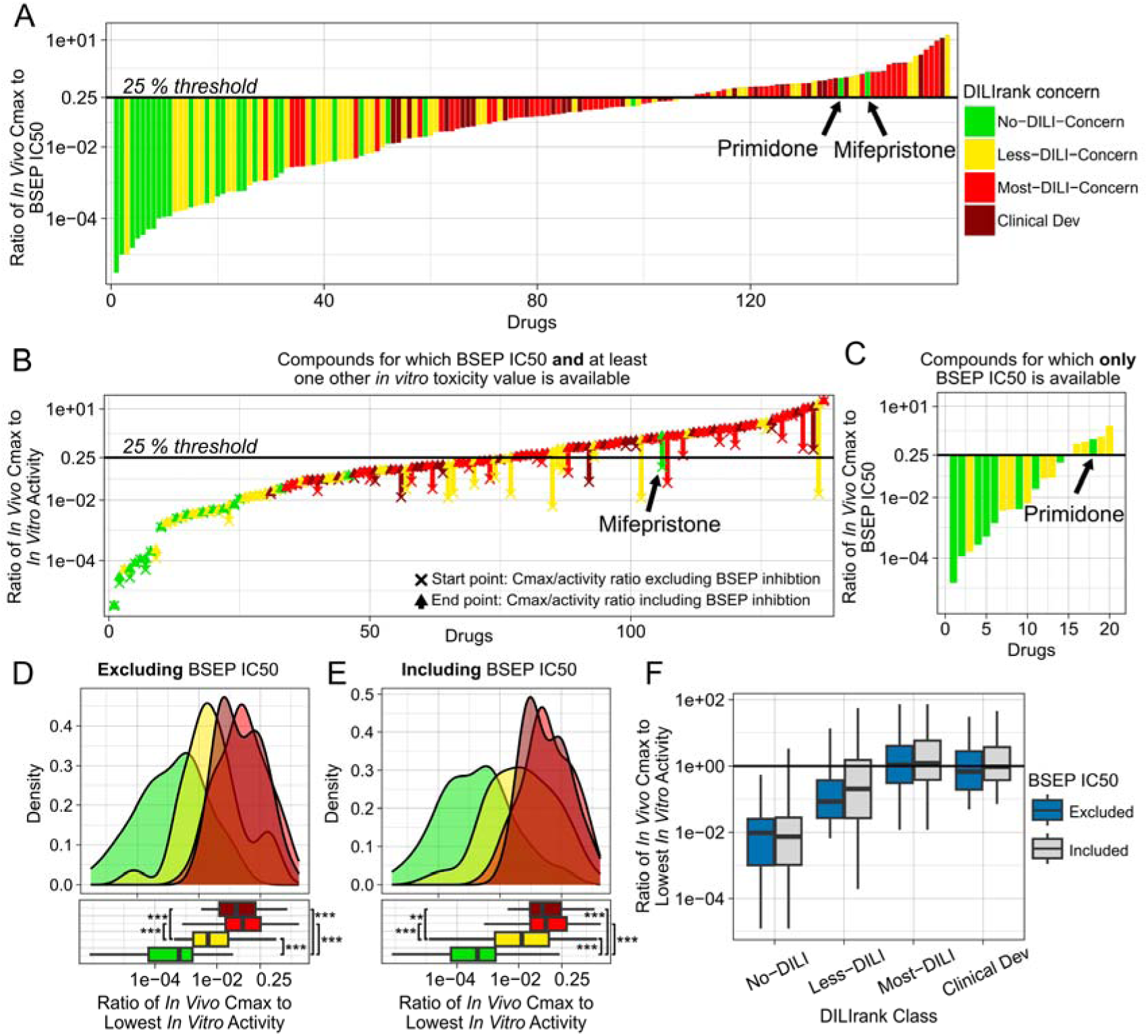
DILI predictivity of *in vitro* BSEP inhibition data (evaluation 4). Ratio of *in vivo* Cmax to BSEP IC50 values of each drug (A). Changes in Cmax to *in vitro* toxicity ratios when including BSEP IC50 values of compounds that have at least one functional *in vitro* toxicity value (B). Ratio of Cmax to BSEP IC50 values of compounds that have no functional *in vitro* toxicity value (C). Probability distribution and boxplot of Cmax to *in vitro* toxicity ratios when BSEP IC50 is excluded (D) or included (E). Boxplot of distributions of Cmax to *in vitro* toxicity ratios of drugs of different DILIrank classes compared with each other when BSEP IC50 is excluded or included (F). Boxplots show the median and interquartile range; whiskers extend to 1.5×IQR. Outliers were excluded from the plot for clarity but were included in statistical analyses. Statistical comparisons on log-transformed values were performed using pairwise Wilcoxon rank-sum tests with Holm correction: * p < 0.05, ** p < 0.01, *** p < 0.001.

Since BSEP inhibition should result in the intrahepatic accumulation of bile acids, one may further expect it to specifically manifest as cholestatic rather than hepatocellular DILI. For this reason, we also tried to collect information about the DILI patterns of drugs. This analysis was more uncertain, because no homogeneously annotated database like DILIrank was available for this. Instead, we collected information about DILI patterns of drugs from various resources. Using our manually curated database, we then also observed that the inclusion of BSEP inhibition most strongly changed the Cmax to toxicity ratios of compounds that were classified as showing a cholestatic rather than hepatocellular DILI pattern (SI-Fig. 2).

Unlike when using functional *in vitro* toxicity data, we observed two drugs that appeared to be false-positive DILI predictions when using BSEP IC50 values. Those were mifepristone and primidone, which are classified as No-DILI concern in DILIrank, yet our analysis indicated that their *in vivo* Cmax values were larger than their BSEP IC50. If those had truly been DILI-negative, then this could potentially have proven that BSEP inhibition on its own may not necessarily lead to DILI. Therefore, we investigated those two compounds more closely.

Mifepristone was developed in the 1980s and is most commonly used for medical termination of pregnancies. In this context, a 200 mg dose is given one single time. We believe this may explain why this drug has not been reported in clinical practice to cause DILI, since such a transient inhibition of BSEP is presumably not sufficiently sustained to cause observable changes in clinical biomarkers. Interestingly, however, we then further found that, since 2012, mifepristone has additionally been approved by the US FDA for the treatment of Cushing’s syndrome. In this context, mifepristone is chronically given to patients once daily. We found three case reports (Funke and Rockey 2019; Shah et al. 2019; Ault et al. 2023) from 2019 to 2023, all published after DILIrank was released in 2016, that present clear cases of Cushing’s syndrome patients in which the treatment with mifepristone was determined to be the cause of their cholestatic DILI. Therefore, we concluded that mifepristone has a DILI risk and that most likely our prediction was not false, but rather that mifepristone’s DILIrank classification is incorrect. In its most frequent use case for abortions, mifepristone may appear safe in clinical practice, but we postulate it can cause cholestatic DILI if it is taken chronically. Likewise, for primidone, we found one case report showing primidone to be the cause of intrahepatic cholestasis in a female patient (Rahimi et al. 2014), as well as a veterinary case reporting liver injury in a dog (Poffenbarger and Hardy 1985). Based on those findings, we considered it more likely that the DILIrank classifications of these two drugs were incorrect, rather than that our mechanistic predictions were.

### Evaluation 5: Dynamic modelling reveals in vivo bile acid perturbations

After performing DILI predictions using static pharmacokinetic (PK; i.e., Cmax) and pharmacodynamic (PD; i.e., IC50) parameters, we evaluated whether integrating PK and PD interactions dynamically would alter our findings. Identical Cmax values can belong to highly different PK profiles, and similarly, DILI also manifests dynamically over time, often requiring prolonged perturbations to become clinically relevant.

To capture those time-dependent effects, we dynamically simulated drug concentrations and consequent *in vivo* inhibition of BSEP within an integrated high-throughput PBK-bile acid (HT-PBK-BA) model. The recently developed bile acid (BA) model (Kister et al. 2025) describes *in vivo* BA dynamics and explicitly represents BSEP as one key transporter involved in BA metabolism. Passing drug concentration profiles predicted by the HT-PBK into the BA model then allows the simultaneous simulation of the dynamic effects of BSEP inhibition and its impact on intrahepatic BA concentrations over time. Given the absence of information on specific drug administration schedules, we assumed a once-daily administration of each drug for one month.

To interpret dynamic HT-PBK-BA simulations, we quantified the duration that intrahepatic BA levels were increased more than two-fold relative to their non-perturbed baseline values or 10% above the non-perturbed peak concentration. When comparing those readouts to static PK/PD predictions based on the ratio of Cmax to BSEP IC50, we found that the results of both approaches correlated well, predicting mostly the same positive and negative responses (Fig. 6CD). However, in some cases, discrepancies emerged (Fig. 6E). For instance, some drugs like troglitazone exhibited rapid decreases in plasma concentrations and therefore caused only negligible intrahepatic BA accumulation in dynamic BA simulations, despite Cmax to BSEP IC50 ratios greater than 1. This showed that the static PK/PD approach may overestimate the clinical toxicity risk of drugs with rapid clearance. Conversely, for other drugs, like bicalutamide, the dynamic BA model indicated relevant perturbations of BA levels, despite them having a low Cmax to BSEP IC50 ratio. This was due to the accumulation of drugs after repeated administrations or slower clearance, which led to relatively prolonged durations of BSEP inhibition.

**Fig. 6:**
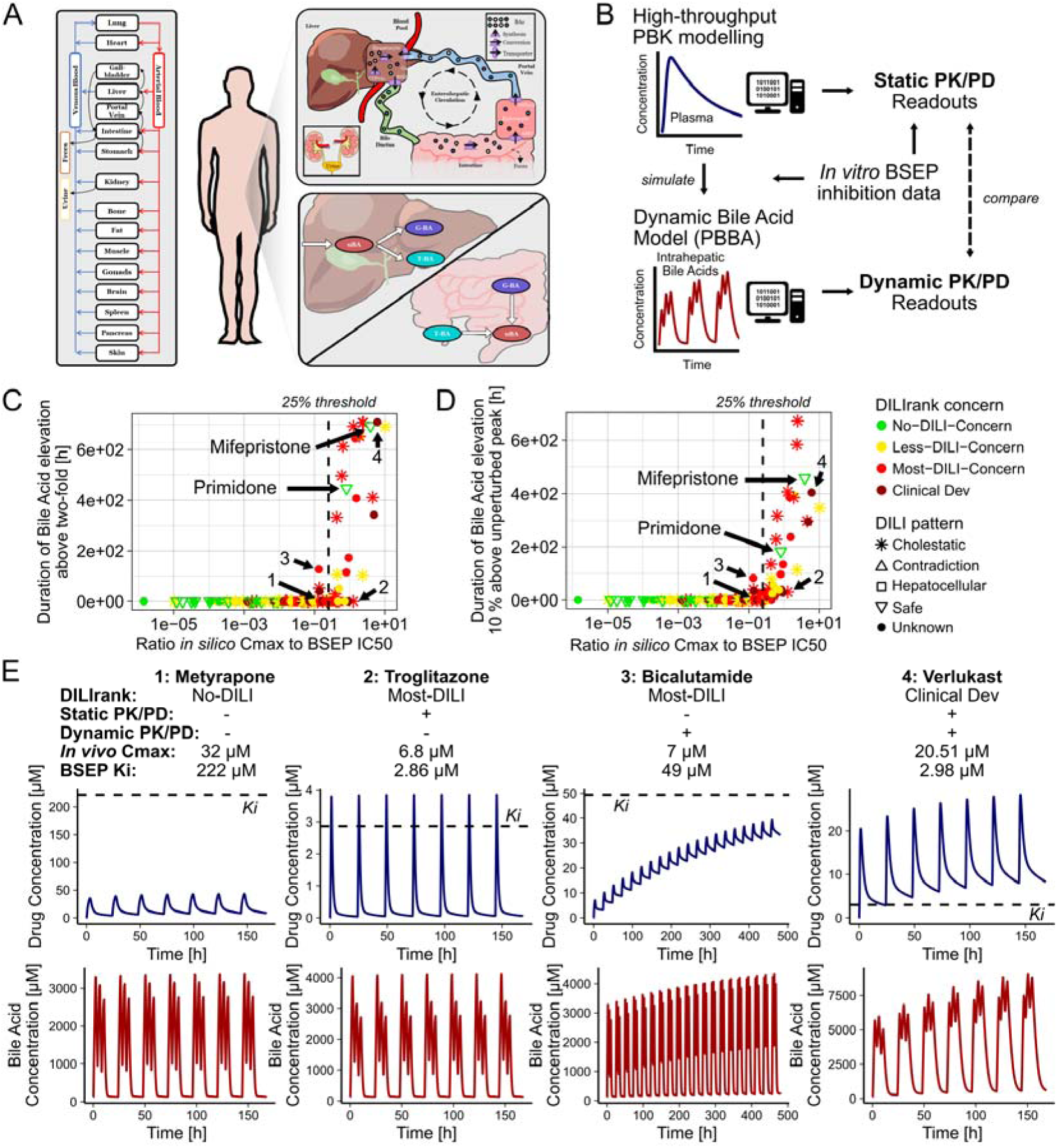
Simulated effects of *in vivo* BSEP inhibition on intrahepatic bile acid levels (evaluation 5). Schematic depiction of processes represented in the Physiologically based bile acid (PBBA) model, graphic is taken from Kister et al. (2025) (A). Schematic representation of the simulation and analysis strategy (B). Dynamic bile acid model perturbations caused through BSEP inhibition by drugs against their ratios of predicted Cmax to BSEP IC50 values and measured by duration during which Bile Acid levels are elevated more than two-fold (C) or duration during which concentrations are more than 10% above physiological peak levels (D). Colour marks DILIrank class and shape the DILI pattern of drugs. Simulated drug and bile acid concentration profiles for four exemplary compounds of which static Cmax to BSEP IC50 ratios and dynamic PBBA model simulation yielded identical or diverging readouts (E). Dashed lines indicate 25% threshold or drug BSEP Ki values.

All compounds showing meaningful BA elevations in dynamic PK/PD simulations were classified as DILI-positive in DILIrank (Fig. 6CD), except for the previously discussed drugs, mifepristone and primidone, which we believe should also be considered DILI-positive. Where information on their DILI pattern was available, all these drugs were also labelled as exhibiting a cholestatic DILI pattern. These results further confirmed our use of 25% as a mechanistically plausible threshold for the Cmax to potency ratio for classifying the expected clinical toxicity outcomes of drugs, as the dynamic BA simulations showed that this value effectively separated drugs that caused meaningful perturbations from those that did not.

Finally, we combined our analyses to identify compounds that may potentially be clinically toxic exclusively through inhibition of BSEP, as such drugs could allow us to gain deeper insights into the clinical effects of BSEP inhibition in humans. We first selected compounds predicted to dynamically cause intrahepatic BA accumulation and subsequently filtered these by only retaining drugs whose *in vivo* Cmax to BSEP IC50 ratios were larger than 0.5 and whose lowest functional potency ratio was lower than 0.5, thus making it unlikely that mechanisms other than BSEP inhibition mediate their DILI risk. This resulted in a list of eight drugs from the original set of 241, which we propose as drugs potentially inducing DILI exclusively via BSEP inhibition (Table 1). With the exception of mifepristone, all of them are classified as DILI-positive in DILIrank, and where information on their DILI pattern was available, it was indicated to be cholestatic. Particularly, pioglitazone and verlukast had *in vivo* Cmax concentrations substantially exceeding their BSEP IC50. Primidone and dicloxacillin also possessed *in vivo* Cmax values larger than their BSEP IC50 but did not possess a functional *in vitro* toxicity value for comparison.

**Table 1:** Compounds causing meaningful intrahepatic bile acid elevations in dynamic model simulations, having Cmax to BSEP IC50 ratios larger than 0.5 and Cmax to functional *in vitro* toxicity ratios lower than 0.5, or no other *in vitro* toxicity value. *Indicates classifications which may be incorrect, as discussed in text. All available values are provided in brackets.

| Compound | DILI rank | DILI pattern | *In Vivo* Cmax | Median BSEP IC50 value(s) | Lowest other, functional *in vitro* toxicity value(s) | Ratio *in vivo* Cmax/BSEP IC50 value | Ratio Cmax/Lowest other, functional *in vitro* toxicity value |
| --- | --- | --- | --- | --- | --- | --- | --- |
| Pioglitazone | Less | Chole-static | 4.18 μM | 0.3 μM (0.098 μM; 0.3 μM; 0.4 μM) | 262.75 μM | 13.94x | 0.016x |
| Verlukast | Clinical Dev | Clinical Dev | 38.8 μM | 3.384 μM | 84.04 μM (84.04 μM; 296.75 μM) | 11.46x | 0.46x |
| Cyclosporine | Most | Chole-static | 0.67 μM | 0.5 μM (0.5 μM; 0.5 μM; 2.393 μM) | 2.3 μM (2.3 μM; 10 μM; 10.1 μM; 14.5 μM; 22.49 μM; 30.8 μM; 39.5 μM; 99.18 μM) | 1.34x | 0.29x |
| Erythromycin | Most | Chole-static | 10.9 μM | 8.55 μM (13.0 μM; 4.1 μM) | 293.7 μM (293.7 μM; 357.6 μM; 740.8 μM) | 1.27x | 0.037x |
| Mifepristone | No* | Safe* | 6.0 μM | 4.71 μM (2.02 μM; 7.4 μM) | 50 μM | 1.27x | 0.12x |
| Fenofibrate | Less | Chole-static | 15.17 μM | 25.39 μM (15.3 μM; 35.48 μM) | 61.07 μM (61.07 μM; 72 μM; 103.18 μM; 103.67 μM; 295.82 μM) | 0.6x | 0.25x |
| Irbesartan | Less | Chole-static | 6.77 μM | 7.31 μM | 580 μM | 0.93x | 0.012x |
| PF-6273340 | Clinical Dev | Clinical Dev | 1.69 μM | 3.0 μM | 42 μM (42 μM; 239 μM) | 0.56x | 0.04x |
| Dicloxacillin | Less | Chole-static | 148 μM | 63.05 μM (56.4 μM; 69.7 μM) | - | 2.36x | - |
| Primidone | No* | Safe* | 41.4 μM | 48.7 μM | - | 0.85x | - |

## Discussion

Here, we demonstrated that high predictivity of the DILI risk of drugs can be achieved by combining *in vitro* hepatotoxicity measurements with pharmacokinetic (PK) information. Simple heuristic predictors used in the pharmaceutical industry, such as drug lipophilicity and dose, showed moderate predictivity, and the integration of PK data with *in vitro* toxicity measurements clearly outperformed them. Notably, exposure-based predictions not only achieved higher predictivity but also aligned with clinically observed DILI severity classifications, highlighting the biological relevance of this approach.

In our analysis, PK information emerged as one of the most influential factors for predicting the DILI risk of drugs. This reflected the narrower distribution of *in vitro* toxicity values compared to Cmax values, making exposure the dominant determinant of clinical toxicity in our dataset. One factor contributing to this phenomenon was likely the exclusion of compounds with extremely low or high potency values outside the concentration test ranges used in *in vitro* assays. Consequently, it remains unclear whether PK information alone would retain such strong predictive power in prospective evaluations of unbiased, early-stage drug candidates. It is plausible that earlier development compounds may exhibit a broader range of *in vitro* toxicity values, given that they are less preselected than the FDA-approved DILIrank drugs evaluated here. But it is also conceivable that toxic potencies of drug candidates are generally biased towards intermediate values. This could be due to the facts that drug candidates are deliberately optimised for bioactivity, making it unlikely for them to be entirely inactive, but also that highly potent toxins presumably only rarely arise spontaneously without active selection. Supporting this hypothesis, toxic potency values within our dataset, even for No-DILI concern drugs, were approximately normally distributed across tested concentrations (Fig. 1J-L). Depending on the distribution of toxic activities of novel drug candidates, PK information may therefore either be considered the dominant or one of two equally critical factors of the clinical DILI risk of drugs. Notably, this is the case despite DILI typically being described to be idiosyncratic without obvious dose dependence.

When combined with Cmax information, *in vitro* BSEP inhibition data not only served as a good predictor of DILI, but it also provided additional information for predicting the risks of drugs whose toxicity would otherwise have been underestimated when solely relying on functional *in vitro* toxicity assays. For example, it enabled the correct prediction of mifepristone’s DILI potential, even though this drug had been classified as No-DILI concern in DILIrank, and for which we only later found strong clinical evidence confirming its DILI potential (Shah et al. 2019; Funke and Rockey 2019; Ault et al. 2023). We further identified several drugs which may cause DILI primarily through BSEP inhibition (Table 1). Alternative toxicity mechanisms, not captured by the functional toxicity assays used here, cannot be excluded for these compounds and hence they need to be evaluated in more detail. But the identification of such drugs would offer a unique opportunity to deepen our mechanistic understanding of the clinical effects of BSEP inhibition in humans.

Notably, compounds whose Cmax to toxicity ratio changed most substantially upon inclusion of BSEP inhibition were classified as Less-DILI concern and exhibited a cholestatic DILI pattern. Similarly, all compounds possessing only BSEP inhibition potential were classified as No- or Less-DILI concern. These findings may indicate that BSEP inhibition alone manifests with lower clinical severity than cytotoxicity or mitochondrial toxicity, but that strong and sustained BSEP inhibition, as observed with mifepristone, can nonetheless lead to significant liver injury. Consistent with earlier studies (Bruijn and Rietjens 2024), our dynamic bile acid (BA) perturbation simulations suggest that most drugs, even those showing a cholestatic DILI pattern, are unlikely to cause meaningful BA level perturbations *in vivo*. This underscores that BSEP inhibition is only one of many mechanisms of drug-induced cholestasis and DILI and emphasises the need to consider the full spectrum of potential toxicity mechanisms.

Similarly, some DILI-positive drugs exhibited Cmax to toxicity potency ratios lower than expected from a mechanistic perspective for causing meaningful toxicity, despite ratios generally correlating well with clinical toxicity outcomes. In the case of estradiol, for example, this discrepancy likely reflected toxicity mechanisms not captured by the *in vitro* assays included here. This highlights the potential to improve DILI prediction accuracies further by incorporating additional endpoints related to other DILI-relevant mechanisms, such as gene expression perturbations, immune responses, or metabolism (Kullak-Ublick et al. 2017).

It is important to note that our analysis could only integrate sparse, heterogeneous data from previously published studies, and not all compounds were tested systematically across all *in vitro* systems. Doing so would likely further improve prediction results. Given that assay sensitivities may differ, applying a single fixed threshold value, such as 25%, may not have been optimal either (Albrecht et al. 2019). Our analysis also did not account for other key factors such as different dosing schedules of drugs, transporters and tissue-specific exposures, nor for possible synergistic effects when multiple toxicity mechanisms co-occur. Integrating such effects, particularly through dynamic simulations as employed for modelling bile acid perturbations, should further enhance clinical toxicity predictions (Watkins 2020). Nevertheless, our findings outline the value of integrating multiple *in vitro* assays, targeting different toxicity mechanisms, as a key factor for the robust predictive performance observed here, which appears to exceed that of most previously published studies (Walker et al. 2020) (SI-Table 1). Given the near-perfect specificity of this mechanistic approach, increasing its sensitivity by incorporating missing influential factors should ultimately enable DILI risk predictions to approach optimal performance using already available *in vitro* and *in silico* methods.

Our analysis also offered an explanation why DILI may typically manifest idiosyncratically in clinical practice, without clear dose dependence. We found that even Most-DILI concern drugs exhibited *in vivo* Cmax concentrations lower than their corresponding *in vitro* potency. Consequently, it is unsurprising that clinically significant toxicity only occurs sporadically, presumably in the most susceptible patient subpopulations, such as those with genetic predispositions, elevated systemic exposures, comorbidities, or comedications. We speculate that a more pronounced dose dependence of idiosyncratic DILI would become observable if drug doses were increased to levels where *in vivo* concentrations consistently exceeded toxic potencies. However, testing this hypothesis in human patients is unethical, and it is likely that maximum recommended doses of drugs were deliberately selected during approval processes to avoid crossing into such clearly toxic exposure ranges.

Here, we focused on the prediction of DILI because liver toxicity remains one of the most significant causes of drug development failures. Unlike other toxicity types, DILI is relatively well-covered by extensive research and some well-annotated resources such as the DILIrank database exist, which supports the evaluation of prediction methods. However, the integrated, mechanism-based approach applied here is not restricted to DILI and can equally be used for other types of drug-induced toxicity, if suitable *in vitro* assays of relevant toxicity mechanisms are available.

Unlike typical machine learning methods that attempt to directly predict DILI risk in a black-box fashion without explicitly representing the underlying toxicity-determining processes, our framework preserves the interpretability and explainability of its results. This enables its evaluation and continuous refinement and also strengthens its potential for regulatory acceptance as an alternative to traditional animal testing. Regulatory agencies worldwide are increasingly committed to using alternative approaches to phase out animal testing, provided that sufficient trust in the reliability of these methods can be established. Our results demonstrate that the required level of reliability, interpretability and predictivity is already achievable today using existing *in*lJ*vitro* and *in*lJ*silico* methods.

## Data availability

All extracted data used for analysis is available in the Supplementary Information and on GitHub (https://github.com/ReneGeci/DILIprediction).

## Code availability

High-throughput PBK simulation code, bile acid model code and analysis scripts are available on GitHub (https://github.com/ReneGeci/DILIprediction).

## Author contributions

Conceptualisation: RG, AZS, LK; Data curation: RG, AZS; Formal Analysis: RG, AZS; Funding acquisition: SS, LK; Investigation: RG, AZS; Methodology: RG, AZS; Supervision: SS, LK; Visualization: RG, AZS; Writing - original draft: RG; Writing - review & editing: AZS, SS, LK

## Abbreviations

AUC: Area under the curve
BA: Bile acid
BSEP: Bile salt export pump
Cmax: Maximum concentration
DILI: Drug-induced liver injury
HT-PBK: High-throughput PBK
LogD: Logarithm of distribution coefficient
LogP: Logarithm of octanol-water partition coefficient
ML: Machine Learning
PBBA: Physiologically based bile acid
PBK: Physiologically based kinetic
PD: Pharmacodynamics
PK: Pharmacokinetics
ROC: Receiver operating characteristic

## Supplementary Material

**Supplementary Table 1:**
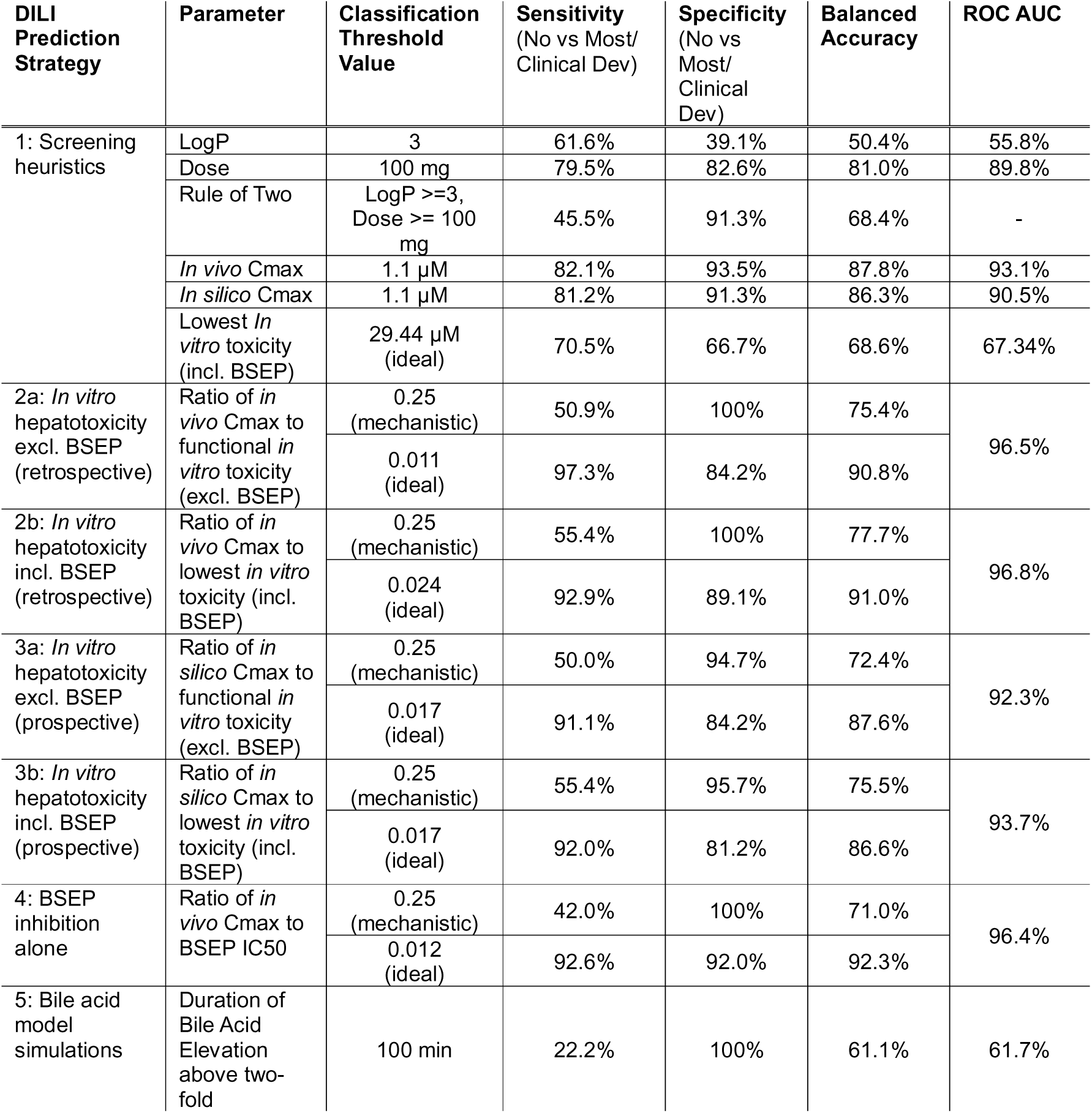
Predictive performance metrics of various DILI prediction strategies. Calculations were based on distinguishing No-from Most-DILI concern and Clinical Development Failure drugs, and assuming that mifepristone and primidone are DILI-positive, despite being classified No-DILI concern in DILIrank, for the reasons outlined in text. Ideal thresholds are based on Youden’s index(YOUDEN 1950).

**SI-Fig. 1:**
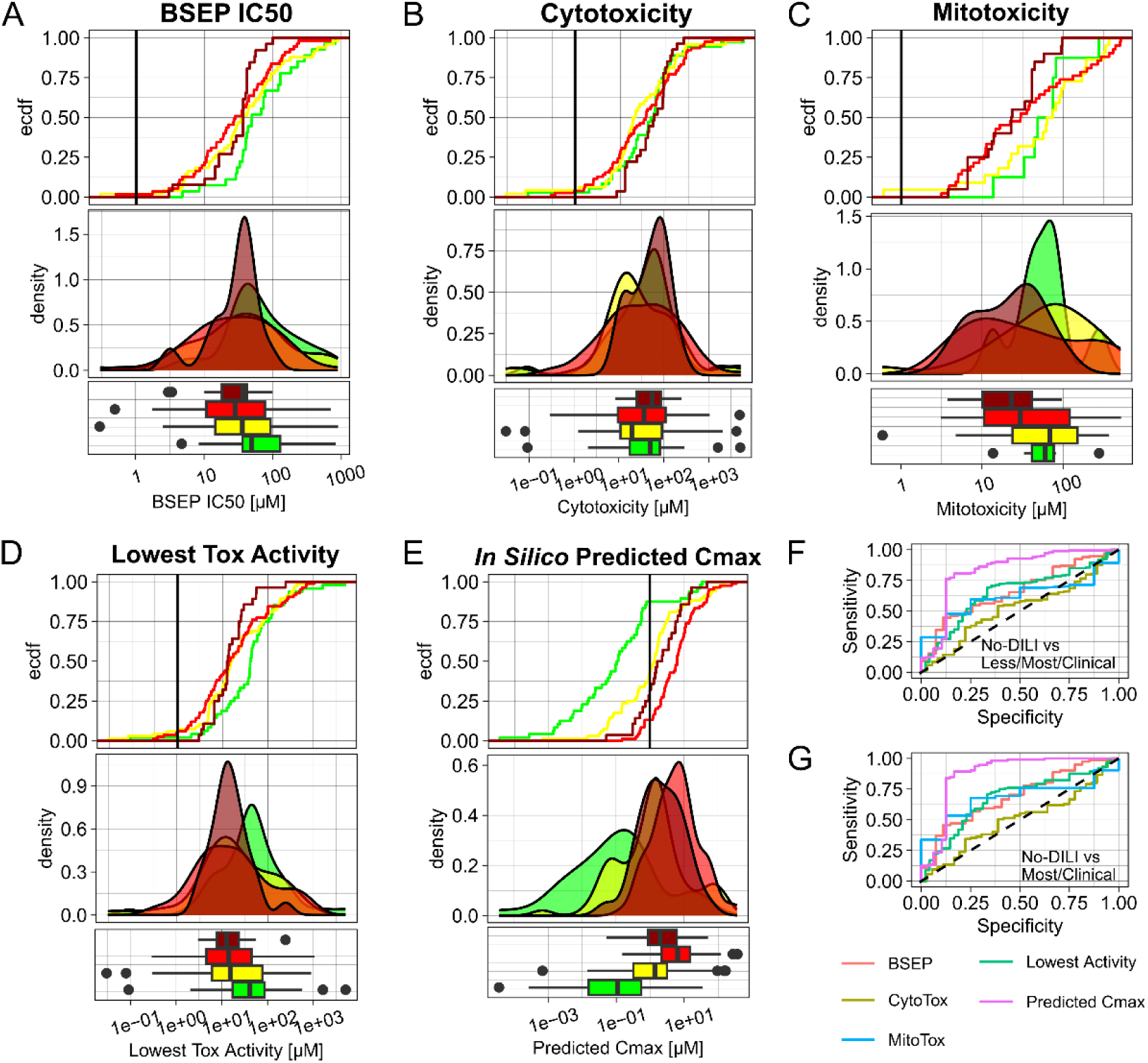
DILI predictivity of i*n vitro* toxicity and *in silico*-predicted Cmax values alone. Cumulative distribution, probability density and boxplots of different properties of drugs of the different DILIrank classes: *In vitro* BSEP IC50 (A), cytotoxicity (B), mitochondrial toxicity (C), the lowest toxicity potency per compound (D) and *in silico*-predicted Cmax values (E). ROC curves of the various parameters for distinguishing No-DILI compounds from Less-/Most-DILI/Clinical Development Failures (F) or only from Most-DILI/Clinical Development Failures. (G).

**SI-Fig. 2:**
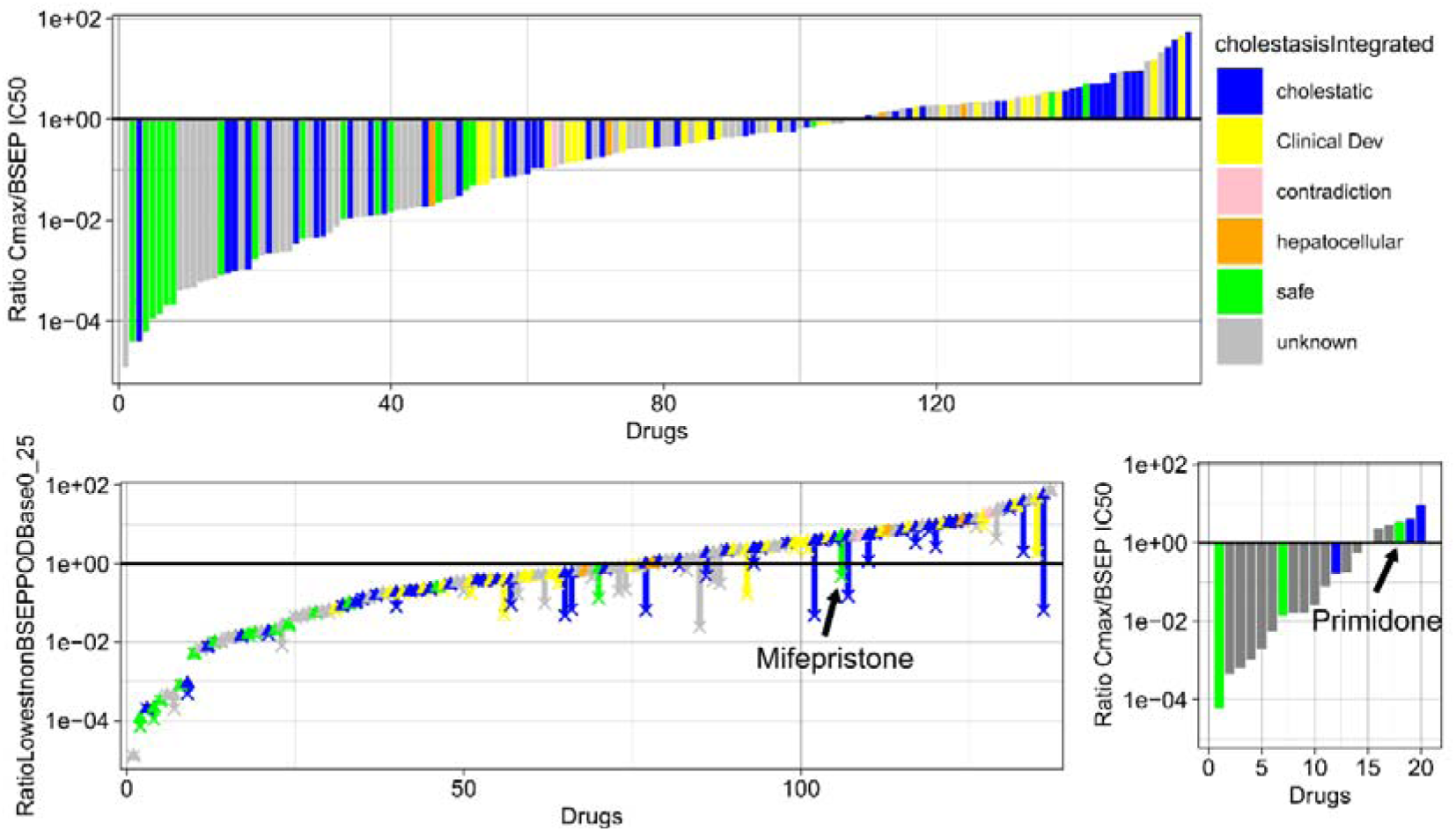
DILI patterns of drugs with BSEP inhibition data. Ratio of *in vivo* Cmax to BSEP IC50 values of each drug (A). Changes in Cmax to toxicity ratios when including BSEP IC50 values for compounds that have at least one functional *in vitro* toxicity values (B). Ratio of Cmax to BSEP IC50 values of compounds that have no functional *in vitro* toxicity value (C).

**SI-Fig. 3:**
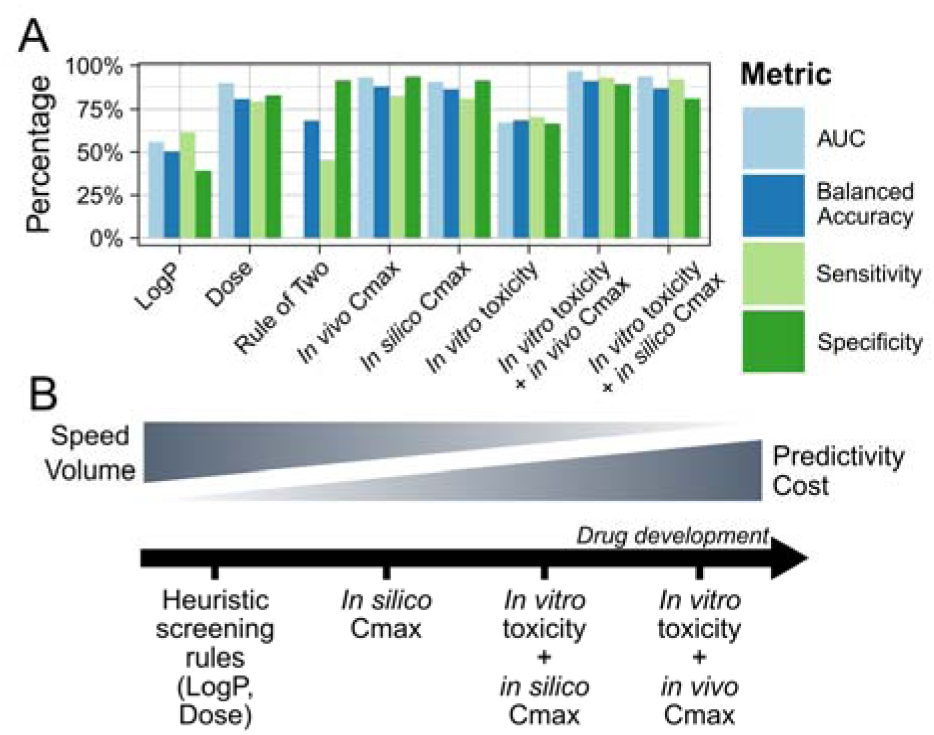
Summary of predictivity evaluations of various DILI prediction strategies. *In vitro* activity refers to the lowest *in vitro* measured toxicity per compound, including cytotoxicity, mitochondrial toxicity and BSEP inhibition (A). Ideal thresholds based on Youden’s index are used. Rule of Two does not possess a ROC AUC value, due to absence of drug risk ranking. Schematic representation of the role of different DILI prediction strategies in the drug development timeline (B).

**SI-Fig. 4:**
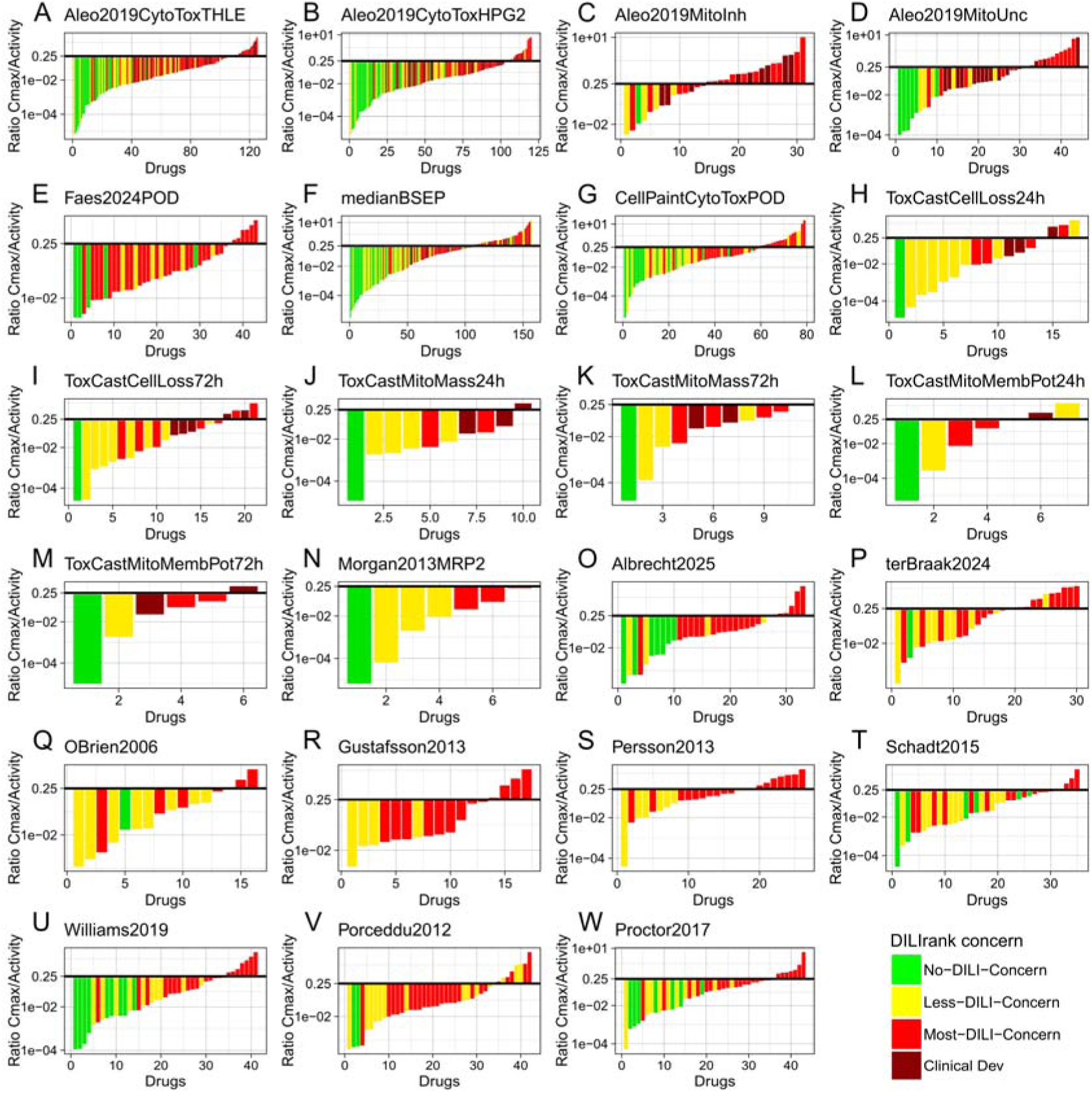
Individual results of each dataset integrated in this study. Ratio of *in vivo* Cmax to lowest functional *in vitro* toxicity of each drug for each *in vitro* toxicity dataset integrated in this study (A-W).

